# Drosophila BEACH domain autophagic adaptor *blue cheese* shuttles between vesicle populations and is required for an early step in autophagy

**DOI:** 10.1101/084434

**Authors:** Joan Sim, Kathleen A. Osborne, Irene Argudo Garcia, Artur S. Matysik, Rachel Kraut

## Abstract

*Drosophila melanogaster blue cheese* (*bchs*) encodes a large BEACH (Beige and Chediak-Higashi) family protein that is postulated to function as an adaptor protein with roles in vesicle trafficking. Mutation in *bchs* leads to the accumulation of ubiquitinated aggregates in aged brains, presumably because of a conserved function with its human homologue Autophagy-Linked FYVE (ALFY), which interacts with Atg5 and p62 to promote the clearance of aggregate-prone proteins. In this study, we present pharmacological and genetic evidence using a well-defined larval motorneuron paradigm that in *Drosophila bchs* mutants, autophagic deficit contributes to neurodegeneration. Specifically, we show that motorneuron death in larvae is accompanied by the accumulation of prominent ubiquitinated aggregates in synaptic termini, and that these are sensitive to autophagy modulating drugs. In primary *bchs* neurons, early autophagic compartments increase in number and intensity based on Atg5 expression, but fail to progress to Atg8-labelled compartments, indicating non-clearance. A rescuing transgene encoding the longest Bchs BEACH domain isoform not only reverses this defect, but also greatly increases Atg8 compartment number and rescues neuronal death. Although only a small fraction of Bchs colocalizes with these markers under wild-type conditions, the population of Bchs that does associate with autophagosomes shuttles between different locations depending on how autophagy is induced. These observations, together with epistatic relationships between *bchs* mutant alleles and autophagy-modulating drugs and genetic backgrounds, points to a model whereby BEACH domain isoforms of Bchs participate in the early steps of autophagy by recruiting Atg5 to target substrates for clearance, and that Bchs’ association with different parts of the autophagy machinery depends upon the type of autophagic stress imposed upon the neuron.

## Introduction

A common pathological hallmark in many neurodegenerative disorders is the accumulation of intracellular toxic aggregates. Degradation of misfolded proteins and protein aggregates is mediated by two main intracellular systems, the ubiquitin-proteasome system (UPS) and autophagy^1,2^. Both pathways are responsible for the recycling of different types of substrates depending on their solubility, half-life and composition, e.g. organelles versus misfolded proteins, type of substrate modification or the presence of specific degradation motifs^3^. Autophagy appears to function in part as a compensatory degradative pathway, because its activity increases when UPS is impaired^4,5^. On the other hand, autophagic efficacy declines with ageing^6^, acting as a possible mechanistic link to sporadic late-onset neurodegeneration.

During macroautophagy, elongation of the isolation membrane requires the E1-like activating enzyme Atg7 and E2-like conjugating enzyme Atg10 to bring Atg12 to Atg5 (an E3-like ligase) which then binds Atg16 to form the Atg12-Atg5-Atg16 multimeric complex^7,8^. This E3-like ligase complex aids in the lipidation of Atg8 protein and dissociates from the membrane upon formation of the autophagosome^9,10^. Although the process of induction by signaling kinases to autophagosome formation and subsequent fusion with lysosomes has been extensively studied^11,12^, the receptors and adaptors that play a role in selective recognition of cargoes in specific cellular locations have been relatively unknown until recently^13^. In mammals one of these adaptors, ALFY (autophagy-linked FYVE protein), scaffolds the machinery associated with isolation membrane elongation around sequestered protein aggregates by binding to the autophagy receptor, p62, through its PH-BEACH (Beige and Chediak-Higashi) domain, Atg5 through its WD40 repeats and phosphatidylinositol 3-phosphate (PI3P)-containing autophagic membrane via its FYVE domain^14,15^.

Loss of function mutations in *Drosophila blue cheese* (Bchs), the orthologue of ALFY, lead to age-dependent accumulation of ubiquitinated inclusions in adult brains, progressive degeneration throughout the nervous system and reduced adult longevity^16^. A targeted genetic modifier screen in which lysosomal and autophagy candidate genes were able to modify a Bchs over-expression induced rough-eye phenotype suggested that Bchs function may be involved in an autolysosomal trafficking pathway^17^. A defect in size and anterograde transport of lysosomal compartments along motorneuron axons in *bchs* mutants supported this premise^18^. In both these studies, *bchs* mutants exhibited morphological features characteristic of atrophying neurons, such as axonal varicosities, ubiquitinated inclusions in the brain and disorganized microtubule bundles. Notably, the degeneration caused by loss of function of *bchs* was never tested for genetic modification by interference with autophagy.

A previous study has reported that ALFY, while not required for autophagy to occur, is associated with autophagic membranes and clears ubiquitinated aggregates from HeLa and neuronal cells^15,19^. However, no studies have examined in detail any changes in the autophagic machinery that may occur in *bchs*, or Bchs dynamic behavior with respect to autophagic components. Therefore, we set out to investigate the spatial localization of Bchs in relation to different steps along the autophagy-lysosomal pathway under various stresses and how these respond to *bchs* mutation.

We first present epistatic arguments, combining loss of function *bchs* alleles with genetic and pharmacological manipulations of autophagy, that Bchs occupies a specific position in an autophagy hierarchy, even though the protein is surprisingly absent from the bulk of autophagic compartments in neurons. Further, we show that rescue of the degeneration by increasing autophagy requires at least one BEACH domain containing member of the three newly identified isoforms, but that this requirement can be bypassed by activating later autophagy events.

## Results

### The *bchs* locus produces three isoforms that are differentially localized with autophagic markers and produce different phenotypes

The existence of a single long isoform of Bchs has been reported, with smaller bands in Westerns being attributed to non-specific cross-reaction of the antibody^17,20^. In our hands, Western blot analysis with a polyclonal antibody raised against the C-terminal 1008 amino acids of Bchs^18^ showed the expression of three Bchs isoforms (approximately 390 kDa, 285 kDa and 250 kDa) both in third instar larval brains and 10-day old adult heads of wild type (WT; *Canton S, yw*) and an *elav* pan-neuronal Gal4 driver (C155, or elav-Gal4) (Fig. 1A,B). The over-expression of Bchs using elav-Gal4 also dramatically increased the levels of all Bchs isoforms in larval brains and heads, supporting the existence of three endogenous Bchs isoforms. Homozygotes for the null allele *bchs17(M)* as well as *bchsLL03462* show negligible expression levels of Bchs (Fig. 1A). As expected from the molecular lesions, the lower Bchs band remains in *bchs58(M)/bchs58(M)* (Fig. 1A, right panel) indicating the presence of the intact second isoform.

**Figure 1.**
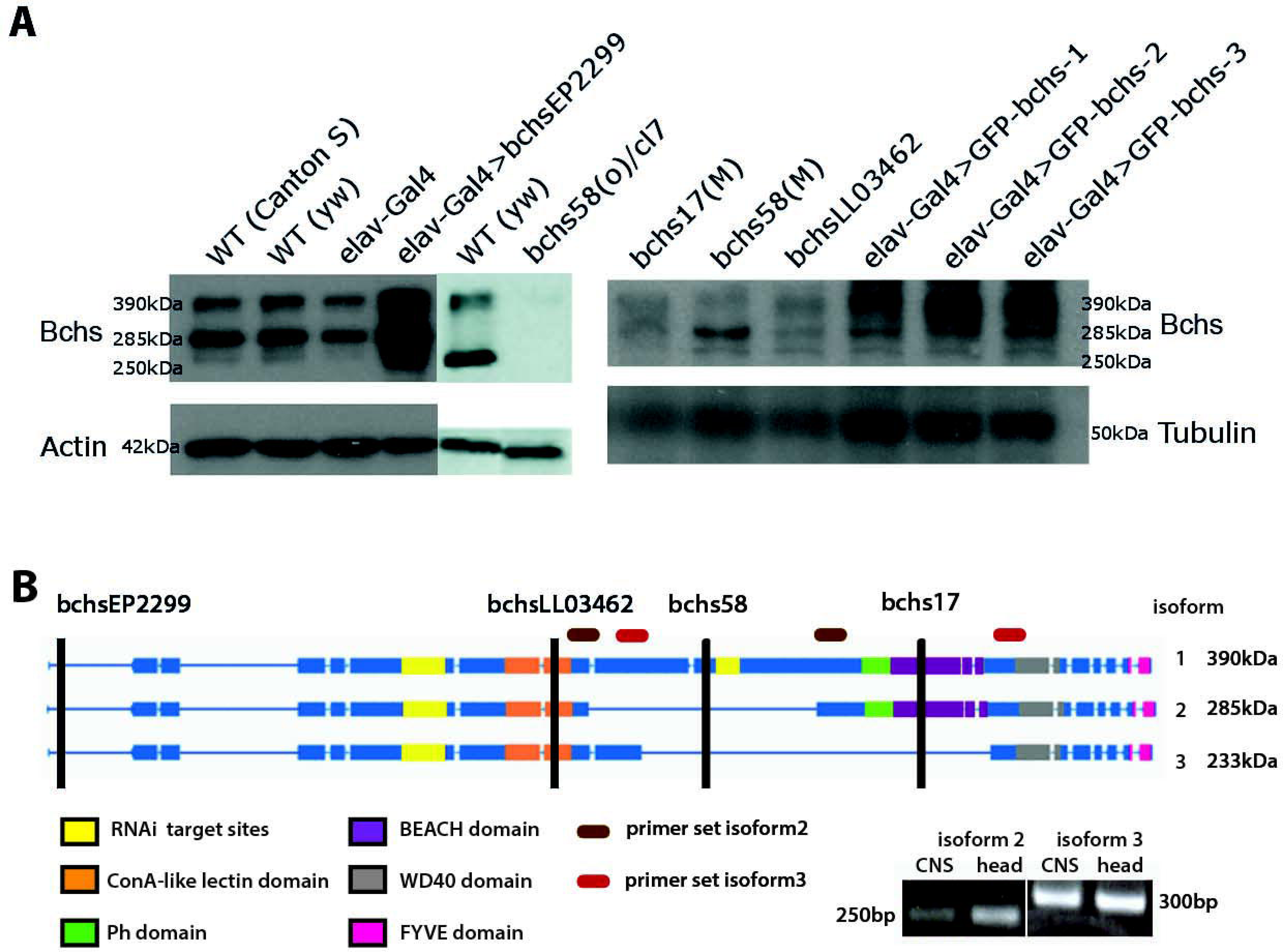
Blue cheese (Bchs) is alternatively spliced. **(A)** Western blot of lysates from third instar larval brains from controls that were wild type (WT) for *bchs* (*Canton S* and *yw*) vs. elav-Gal4-driven overexpression of the endogenous gene via EP2299 (elav-Gal4>bchsEP2299), from different *bchs* mutations (*bchs58(O), 17(M)*, and *>LL03462* >alleles), and from elav-Gal4 overexpression of the three isoforms (elav-Gal4>GFP-bchs1,2,3). Actin and tubulin were used as loading controls in different experiments. **(B)** Schematic illustrating domains excluded from identified splice isoforms identified by RT PCR from larval CNS and adult head. The positions of *bchsEP2299* insertion, *bchsLL03462, bchs58, bchs17* mutations are indicated with vertical black lines. Splice isoforms 1, 2, and 3 have expected sizes of 390kDa, 285kDa and 233kDa, respectively. Using the primer sets shown to detect splice isoform 2 (brown ovals) and splice isoform 3 (red ovals), 250bp and 300bp bands were detected, from which we deduced that splice isoform 1 is 2.8kb larger than isoform 2 and 4.1kb larger than isoform 3.

The three isoforms of Bchs (depicted in Fig. 1B) were cloned as N-terminal GFP fusions and expressed in larval brain via elav-Gal4, and these were dissociated into primary neuronal cultures. Live imaging of these neurons in the background of transgenic RFP-Atg5 and mCherry-Atg8a expression to label early and late autophagic compartments showed that where GFP-Bchs can be detected in vesicles, only BEACH-containing isoform 1 (GFP-bchs-1) is predominantly coincident with autophagosomes (Fig. 2A,B), whereas isoforms 2 and 3 have much less association with either Atg5 or Atg8 (Fig. 2C-F), and indeed are often mutually exclusive (this can be seen particularly well in Fig. 2D and F). Autophagosomal association does not appear to depend on the BEACH domain, since BEACH domain-containing isoform 2 is less autophagosomal than isoform 3, which has none.

**Figure 2.**
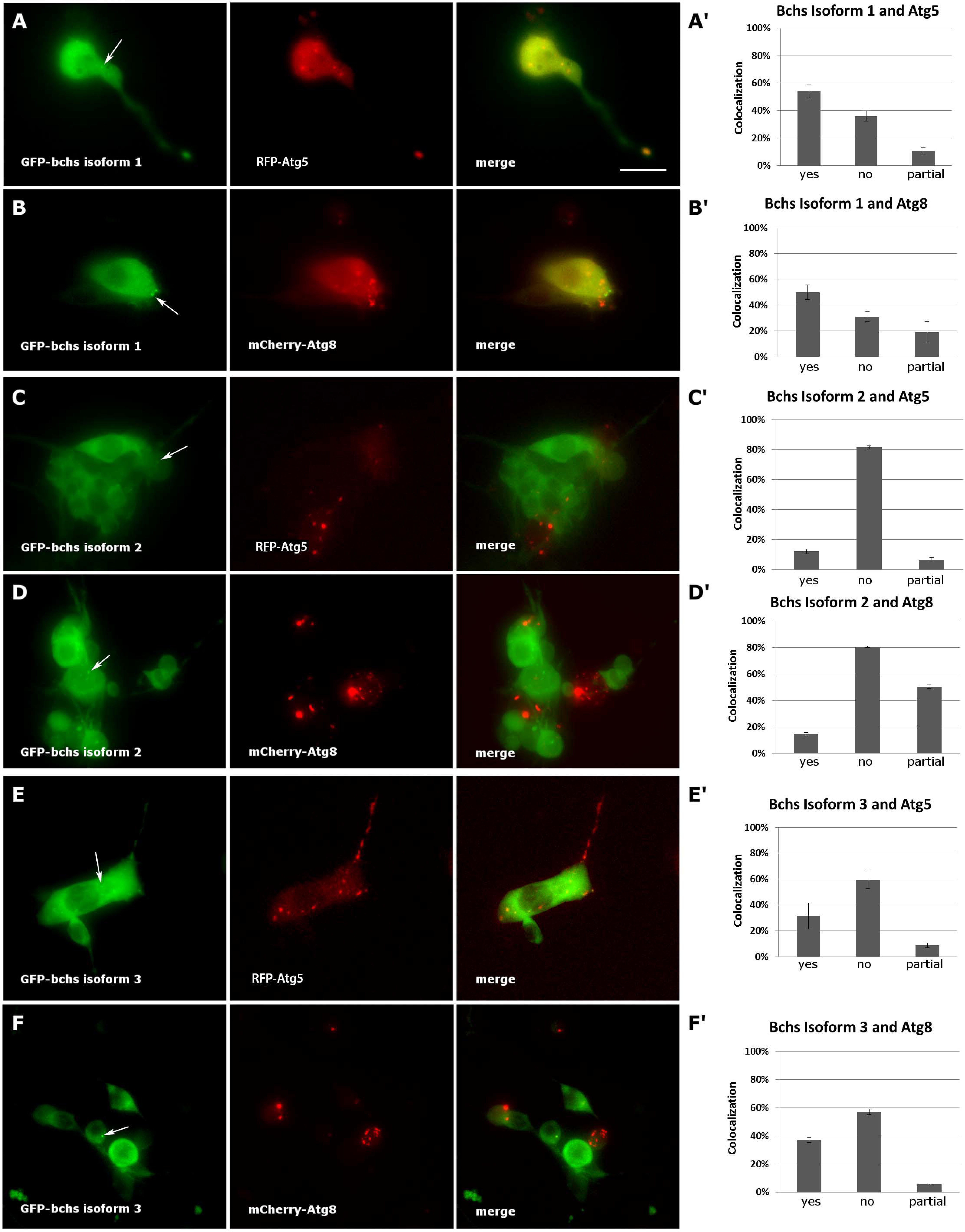
Blue cheese isoforms colocalize to different extents with autophagosomes. Live images and colocalization tally of elav-Gal4-driven **(A, A’)** GFP-bchs-1, the longest isoform, with RFP-Atg5 in primary larval neurons; **(B, B’)** GFP-bchs-1 with mCherry-Atg8a; **(C, C’)** GFP-bchs-2, which retains a BEACH domain, with RFP-Atg5; **(D, D’)** GFP-Bchs-2 with mCherry-Atg8a; **(E, E’)** GFP-Bchs-3, the non-BEACH isoform, with RFP-Atg5; **(F, F’)** GFP-bchs-3 with mCherry-Atg8a. Arrows point to vesicularly located GFP-bchs isoforms. Scalebar = 10 μm.

### Autophagic modulation of neurodegeneration in loss of function *bchs* mutants depends on the BEACH domain

A specific function of Bchs in aggrephagy was strongly suggested by the demonstration that Bchs and its homologue ALFY interact physically with the autophagy machinery, and that both are able to reduce aggregates induced in human and *Drosophila* cells^15^. However, a loss of function phenotype in the fly was not tested for interactions with either pharmacological or genetic modulators of autophagy, which should occur if *Drosophila* Bchs functions as an autophagic adaptor. As in our earlier study^18^, the loss of larval motorneurons aCC and RP2 was used as a means of representatively quantifying the extent of neurodegeneration in *bchs* alleles that differentially affect the three isoforms, and to assess the effects of autophagy modulating drugs in these genetic backgrounds. These two neurons, which were shown to have a high frequency of TUNEL-labelling in the previous study, are specifically labelled in our assay by driving membrane-localized CD8-GFP with an *even-skipped* driver (this driver-reporter combination, used in Figs. 3-5, is referred to as eve>GFP).

*bchs58(O)*, isolated by Khodosh *et al* as *beach58* by EMS mutagenesis in the EP2299 UAS line, was characterized as a strong loss of function mutation^16,18,20^. Our sequence analysis of this allele detects an insertion of 3 bases and a deletion of 17 bases, resulting in a frame shift and subsequent stop codon. Thus, the *bchs58* lesion should remove only the longest of the three Bchs splice isoforms identified in the present study but leave the two shorter isoforms intact, as Fig. 1A suggests. *bchs58(M)* was generated by precise excision of the EP element from *bchs58(O)*, and exhibits a lower penetrance of motorneuron death than *bchs58(O)* (Fig. 3A). Similarly, *bchs17(M)* is a precise excision of the EP2299 element from *bchs17(O)*, which introduces a stop codon at Trp2640. *bchs17(M)* has a higher penetrance of motorneuron death than *bchs58(M)* (Fig. 3A), consistent with its removing both BEACH containing isoforms (Fig. 1A,B).

**Figure 3.**
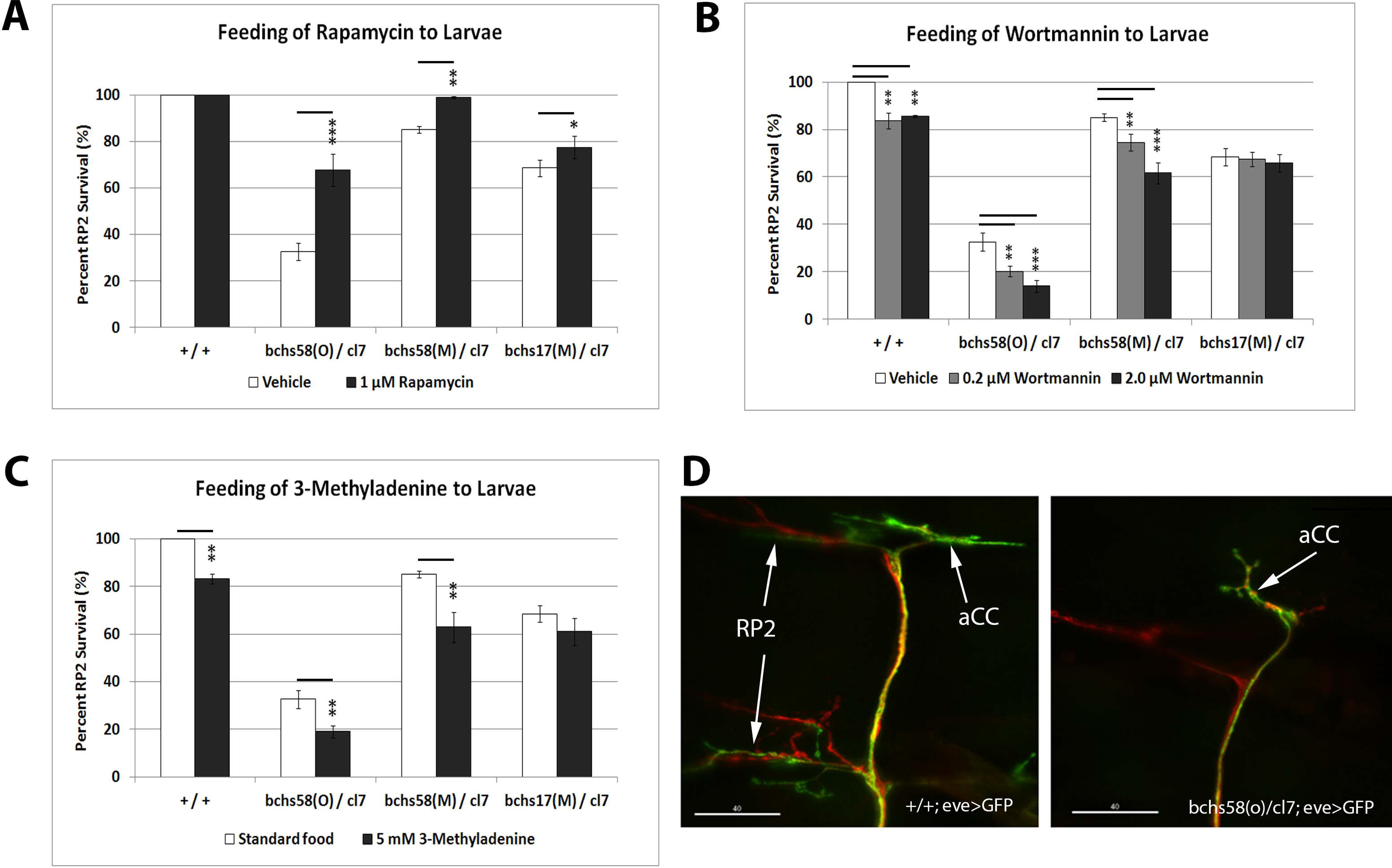
*bchs’* neurodegenerative phenotype is sensitive to autophagy-modulating drugs fed to larvae. The indicated wild-type control and *bchs* mutant genotypes were fed on standard food containing either **(A)** rapamycin, **(B)** Wortmannin or **(C)** 3-methyladenine. All third instar larval motorneurons were labelled with anti-Futsch (22C10; red), and co-labelled with anti-GFP to detect aCC and RP2 motorneurons (green; arrows), examples of which are shown in **(D)**. RP2 is absent in the *bchs* mutant (right-hand panel). Percent survival of RP2 motorneurons over ~100 hemisegments was calculated for each treatment, and performed in triplicate. Scalebar = 40 μm. Chi-square test was used to determine statistical significance; * *p* < 0.05, ** *p* < 0.01, *** *p* < 0.001.

We used the motorneuron death assay to test the effects of rapamycin, a well-established autophagy inducer whose mode of action involves inhibiting the target of rapamycin complex 1 (TORC1), thereby alleviating the suppression of autophagy^21,22^. After rapamycin feeding, both the strong loss of function allele over a deficiency, *bchs58(O)/cl7*, and the hypomorphic excision allele over deficiency, *bchs58(M)/cl7*, showed a significant amelioration of RP2 death, while *bchs17(M)/cl7* showed a lesser effect (Fig. 3A). This suggests that autophagy is deficient in *bchs* mutants and can be substantially compensated through rapamycin feeding, but only if the BEACH-containing 2^nd^ isoform is unaffected by the mutation, as is the case in *bchs58. bchs17(M)*, which should result in a truncation within the BEACH domain of the two longer isoforms, was refractory to the effects of rapamycin, compared to *bchs58(O)* (Fig. 3A). As rapamycin triggers autophagy by inhibition of TORC1 and subsequent activation of the Atg1 complex, an epistatic interpretation of this result suggests that Bchs acts downstream of Atg1 in the autophagy hierarchy. It also identifies BEACH-containing isoforms of Bchs as the primary participants in autophagy.

Wortmannin and 3-methyladenine (3-MA) are widely used broad-spectrum phosphatidylinositol 3-kinase (PI3K) inhibitors that act mainly via the suppression of class III PI3K activity to inhibit the nucleation of phagophores through PI3P production^23,24^. Grown in 0.2 µM Wortmannin, wild-type control, *bchs58(O)/cl7* and *bchs58(M)/cl7* showed a significant reduction of motorneuron survival; this is further reduced by a ten-fold increase of Wortmannin concentration in *bchs* mutants (Fig. 3B). In contrast to the *bchs58* alleles, Wortmannin did not exacerbate *bchs17(M)/cl7* (Fig. 3B), a genetic background that lacks the two BEACH-containing isoforms. Similarly, 3-MA caused motorneuron death in wild-type (phenocopying *bchs*) and exacerbated *bchs58(O)/cl7* and *bchs58(M)/cl7*, but did not significantly exacerbate *bchs17(M)/cl7* (Fig. 3C). These data again point to the BEACH domain isoforms being responsible for the residual autophagic capacity that appears to be present in the rescuable *bchs58* alleles.

### Activation of a late-stage autophagy step by *atg7* rescues the *bchs* degenerative phenotype

There is evidence that ALFY interacts physically with the autophagy machinery, but a *bona fide* genetic interaction between a loss of function *bchs* phenotype and autophagy has not been tested. Therefore, we wanted to examine whether the manipulation of autophagy genes could modify *bchs* motorneuron death. Atg7, an E1-like ubiquitin activating enzyme that is involved in the conjugation of phosphatidylethanolamine (PE) to Atg8, has been shown to promote neuronal health by suppressing the accumulation of ubiquitinated aggregates, thereby contributing to *Drosophila* adult longevity^25,26^. RT-PCR of mRNA isolated from adult heads of *Atg7[EY10058]* which has upstream activating sequences (UAS) inserted before the *atg7* transcriptional start site driven with elav-Gal4 (labelled elav-Gal4>atg7 in Fig. 4A), showed an increased level of *atg7* transcripts (Fig. 4A, lane 3). Therefore, we used this line to over-express Atg7 in the presence or absence of Bchs.

**Figure 4.**
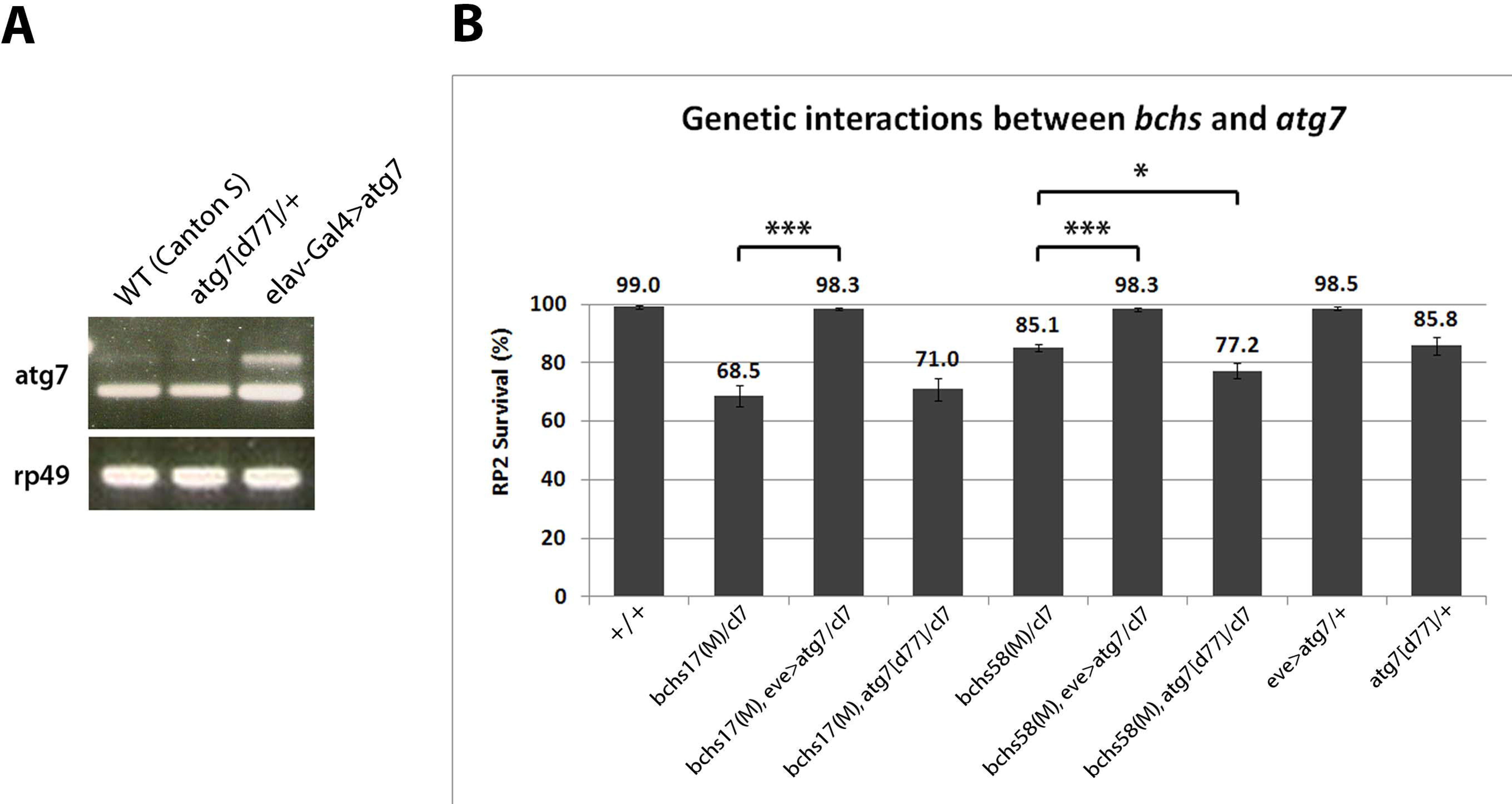
Genetic enhancement of autophagy rescues *bchs* neuronal death. Over-expression of *atg7* ameliorates neuronal death in *bchs* mutants while a single allelic deletion of *atg7* does not have an effect on *bchs* mutants. **(A)** Reverse-transcription PCR on total RNA from adult heads of wild type (WT, CantonS), atg7[d77]/+ heterozygotes, and elav-Gal4>atg7, using primers specific for *atg7* mRNA and *rp49* rRNA as loading control. **(B)** Percentage survival of RP2 motorneurons in third instar larvae of the indicated genotypes, based on presence of GFP in the RP2 motorneuron; experiments performed in triplicate. All genotypes include the driver/reporter combination eve>GFP in homozygosity. Error bars represent standard deviation, and Chi-square test was used to calculate statistical significance. * *p* < 0.05, ** *p* < 0.01, *** *p* < 0.001.

Strikingly, over-expression of Atg7 via eve-Gal4 (labelled eve>atg7 in Fig. 4B) rescued motorneuron survival to almost 100% in both *bchs17(M)/cl7* and *bchs58(M)/cl7* (Fig. 4B). Notably, this genetic augmentation of a later-occurring autophagy step was more efficacious than rapamycin feeding, which was only able to very mildly improve *bchs17(M)/cl7* degeneration (Fig. 3B). *bchs58(M)/cl7* showed a similar percentage of neuronal death as the loss-of-function allele *atg7[d77]/*+ (85.1% vs. 85.8%) and combining them resulted in a further reduction of neuronal survival from 85.1% to 77.2%. This was not the case for *bchs17(M)/cl7*, where combination with *atg7* heterozygosity did not significantly exacerbate the phenotype (68.5% vs. 71%) (Fig. 4B). Another *atg7* deletion *atg7[d14]* was synthetic lethal when recombined with *cl7*, making it difficult to examine the effect of *atg7-* homozygosity on *bchs*.

Similarly to what we saw with Wortmannin and 3-MA (Fig. 3C,D), *bchs58(M)*, where BEACH-containing isoform 2 is intact, is hypomorphic, whereas *bchs17(M)*, which removes both BEACH-containing isoforms, behaves like a loss of function amorph which cannot be further exacerbated by disrupting autophagic activity. Furthermore, the epistatic rescue of both hypomorphic and strong *bchs* alleles by *atg7* over-expression suggests that *atg7* and *bchs* act in the same pathway, with *atg7* acting downstream of *bchs*.

Atg1 activity has previously been shown to rescue phenotypes that result from autophagic deficit by overexpression of a transgene^27^. We attempted this in the background of *bchs* alleles, using two different available constructs (UAS-Atg1^GS10797^, and UAS-Atg1(6A), a kind gift of T. Neufeld), but both of these resulted in nearly complete motorneuron death when expressed in the neurons to be assayed, both alone and in combination with *bchs* alleles, and could therefore not be tested for rescue.

The functionality of Bchs-GFP isoform 1—the most autophagosome-associated isoform—was assayed for its ability to rescue the motorneuron loss of different *bchs* loss-of-function alleles. To test this, GFP-bchs-1 was recombined onto the *cl7* deficiency chromosome, and crossed into the background of different *bchs* combinations. *bchsLL03462*, a strong allele resulting from an insertion into *bchs* of a splice acceptor site followed by stop codons into the 7^th^ coding exon at aa 1229 preceding the BEACH domain (Flybase allele report FBti0124589) by itself gave only ~40% motorneuron survival, but was rescued by the transgene GFP-bchs-1 to ~100% survival (Fig. 5A). The ~68% survival seen in *bchs17(M)* was mildly rescued to ~80%, while *bchs58(M)* was not rescued (not shown).

**Figure 5.**
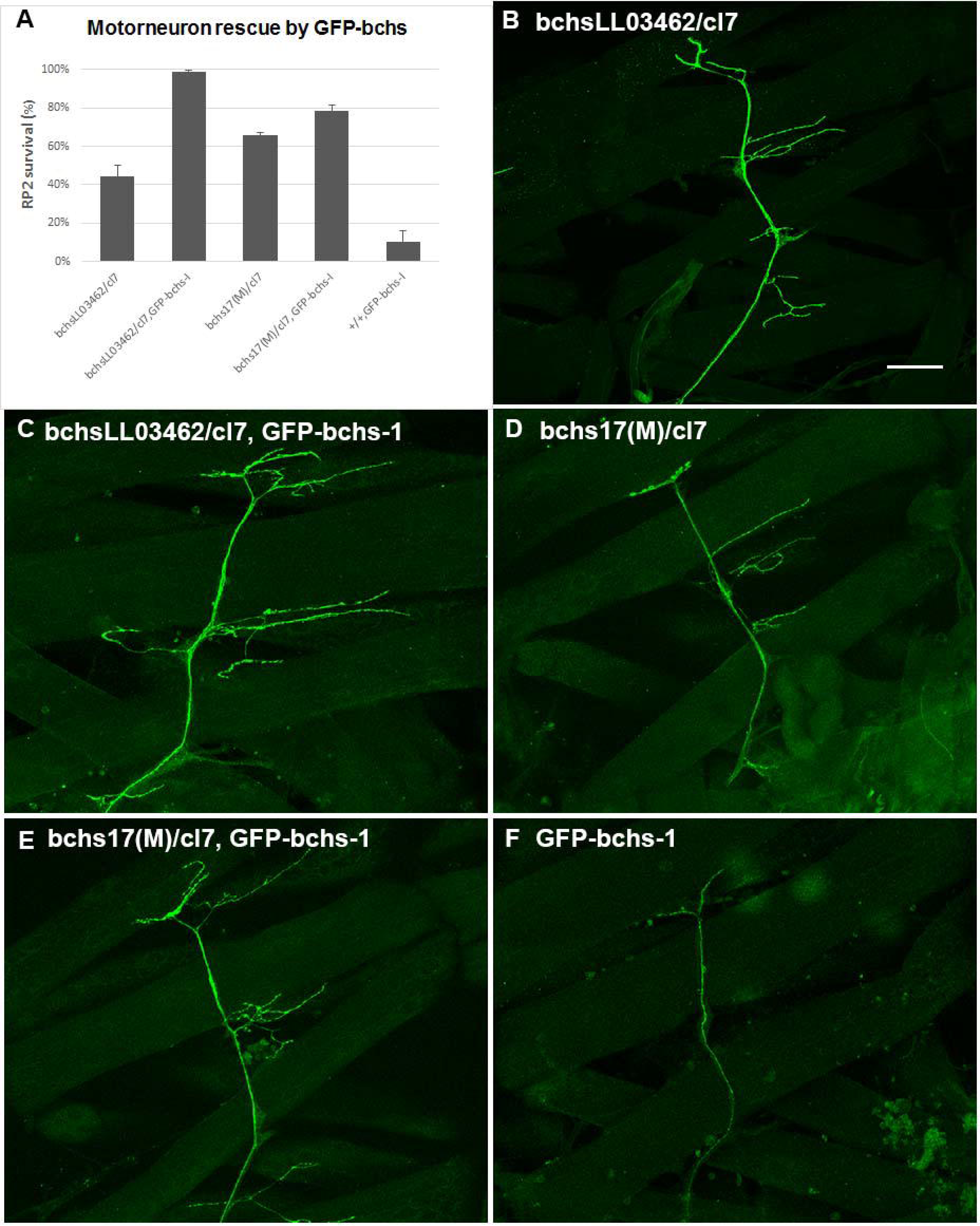
Expression of full length Bchs isoform 1 (GFP-bchs-1) rescues the larval NMJ phenotype and neuronal death of *bchs* loss of function, but kills motorneurons by itself. **(A)** Quantification of RP2 motorneuron survival after GFP-bchs-1 expression via eve-Gal4 in *bchs* mutant backgrounds *bchsLL03462/cl7, bchs17(M)/cl7*, or in the control background (+/+). **(B-F)** images of representative NMJs in the larval body wall labelled with eve>GFP in the same *bchs* backgrounds, with or without GFP-bchs-1. Scalebar =10 µm.

### Accumulation of ubiquitinated aggregates in *bchs* motorneuron termini accompanies degeneration and is abolished by augmenting autophagy

Previous reports have demonstrated the accumulation of insoluble ubiquitinated protein aggregates in aged *Drosophila bchs* adult heads and in photoreceptor axons of Bchs over-expressing late pupae^16,17^. Larval motorneurons are ideally suited to examine the subcellular localization of such aggregates due to their clearly identifiable anatomy. Therefore, we investigated whether ubiquitinated aggregates accumulate in *bchs* motorneurons and where this occurs.

Aggregates indeed appear prominently in the dorsal-most synaptic arbors of motorneurons in third instar larval body walls of *bchs* animals (Fig. 6A). There was no obvious accumulation of ubiquitinated conjugates in motorneuron axons and cell bodies in *bchs* mutants (data not shown). Aggregate area sizes were categorized into three groups: 0 - 1 µm^2^ (small), 1.1-10 µm^2^ (medium) and 10.1 - 50 µm^2^ (large). We observed that while the smallest aggregates were nearly always found in wild type termini, the frequency of intermediate- and large-sized ubiquitinated aggregates was much higher in the various *bchs* mutants. The null allele *bchsLL03462/cl7* had the highest occurrence of intermediate aggregates, whereas *bchs58(O)/cl7* had fewer overall (Fig. 6B). Although unexpected based on the severity of the alleles, this may be related to the possibly neomorphic nature of the 58 allele.

**Figure 6.**
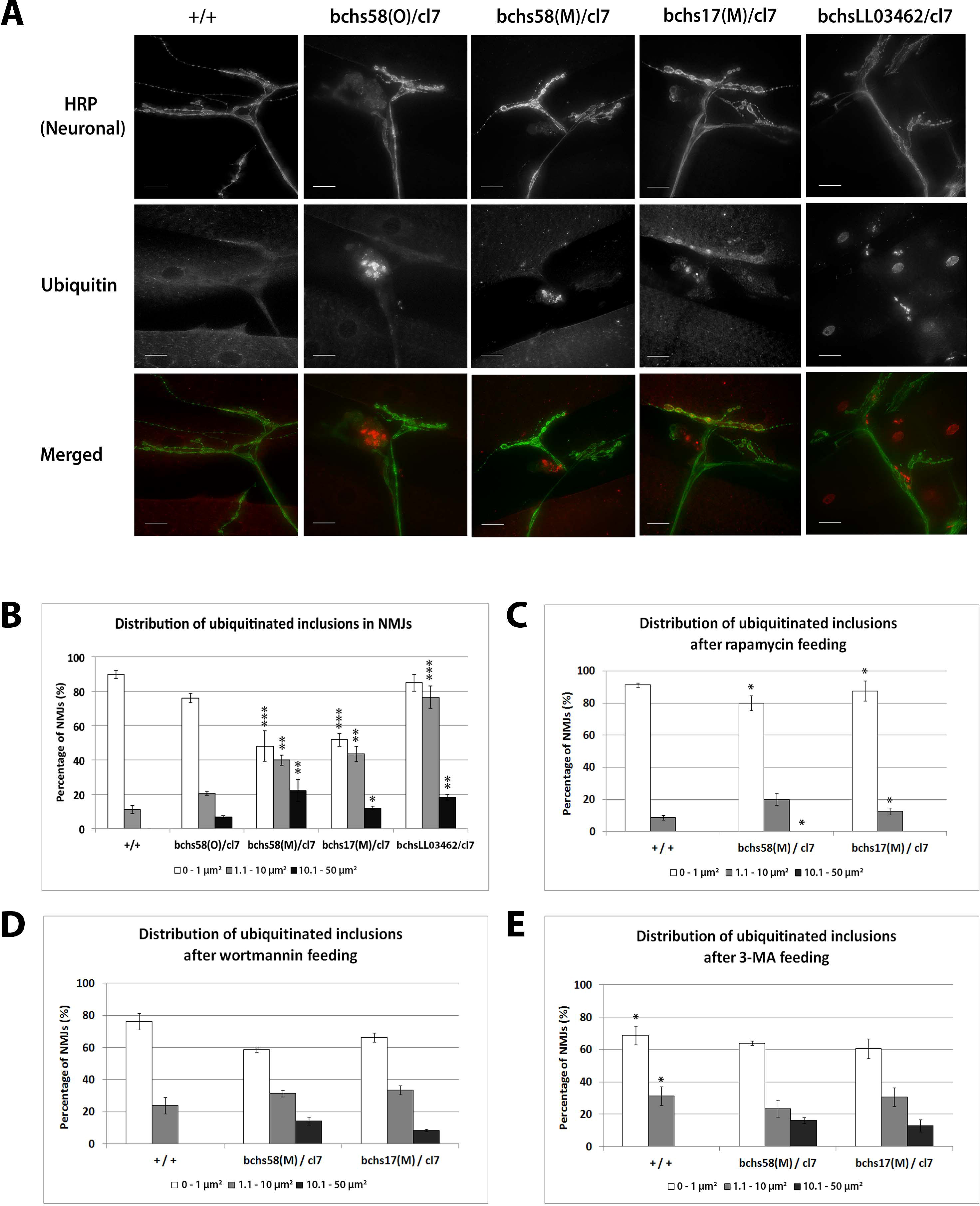
Ubiquitinated aggregates accumulate in neuronal termini of *bchs* larval NMJs and can be reduced by rapamycin feeding. **(A)** Third instar larvae of wild-type control and *bchs* mutants were dissected and immunostained for anti-HRP (a Drosophila pan-neuronal marker) and anti-poly-ubiquitin. The scale-bar represents 20 µm. **(B)** The distribution of the size of ubiquitinated inclusions was analyzed using ImageJ and categorized into three groups: 0 - 1 µm^2^, 1.1 - 10 µm^2^ and 10.1 - 50 µm^2^. The percentages of NMJs with ubiquitinated inclusions in each of the three groups were calculated for the different genotypes which were fed on standard food and performed in duplicates. The error bars represent S.E.M. Chi-square test was used to determine pairwise statistical significance between frequency of occurrence of a given aggregate class in *bchs* mutants vs. the same class in the wild-type control. * p < 0.05, ** p < 0.01, *** p <0.001. Larvae of the indicated genotypes were fed on standard food containing either **(C)** rapamycin, **(D)** Wortmannin or **(E)** 3-methyladenine from first instar to third instar before repeating the same procedures for analysis. Stars in C and E show significance for pairwise comparisons of the distribution of ubiquitinated inclusions with the same genotype in **B** without drug.

We next investigated whether autophagy-modulating drugs affected the distribution of aggregates in termini, and whether this correlated with motorneuron viability. For this purpose, rapamycin, an enhancer of autophagosome formation and flux^28,29^, was fed to larvae during their development. *bchs58(M)/cl7* and *bchs17(M)/cl7* fed 1 µM rapamycin showed significantly lower percentages of neuromuscular junctions (NMJs) containing medium and large ubiquitinated aggregates, while small ubiquitinated aggregates increased significantly in number (Fig. 6C). We conclude that rapamycin increases motorneuron survival in *bchs* mutants at least in part by promoting the degradative removal of ubiquitinated aggregates in synaptic termini.

In contrast to rapamycin feeding, Wortmannin and 3-MA did not affect the distribution of ubiquitinated aggregates in *bchs* mutants (Fig. 6D and 6E). However, 3-MA feeding caused a significant shift towards the formation of medium (but absence of large) ubiquitinated aggregates in wild-type larvae (Fig. 6E). In order to demonstrate that rapamycin enhanced autophagy in our system, we examined muscles in rapamycin-fed wild type larvae. We found that while Wortmannin feeding suppressed puncta formation in the mCherry-Atg8a marker driven by the muscle driver 24B, rapamycin led to an increase in punctae, suggesting that these drugs have the expected effects (Supplementary Fig. 1).

### Early autophagosomes increase in *bchs* primary neurons and Bchs expression drives progression to late autophagy

Since the observed accumulation of ubiquitinated aggregates in neuronal termini of *bchs* NMJs suggests a blockage in the autophagy pathway, it was of interest to investigate in which step this blockage might be taking place. Therefore, the subcellular localization of Atg5, which marks the pre-autophagosome (but not completed autophagosome) and Atg8, which marks the mature autophagosome and autolysosome, were examined. No difference was detectable in the deposition of Atg5 or Atg8 punctae along axons and synaptic termini of aCC and RP2 motor neurons between wild-type control and *bchs* mutant larvae (data not shown).

Primary neurons were cultured from third instar larval brains in order to obtain a broader spectrum of neuronal types than just motorneurons, and because these neurons could be imaged and analyzed in much larger numbers in order to obtain better statistics. We assessed the number, size, and brightness of Atg5- and Atg8-positive compartments in primary neurons prepared from 3^rd^ instar larvae, labelled with antibody vs. Atg5 or Atg8 and anti-poly-Ubiquitin (Fig. 7). Atg5-positive compartments accumulated throughout the cell and increased in number and/or brightness in all *bchs* allelic combinations (Fig. 7A-D), whereas Atg8-positive compartments were largely unchanged (Fig. 7E-H) (see also Supplementary Fig. 2). Ubiquitin-labelled inclusions were visible in all preparations, but appeared more prominent and vesicular rather than predominantly nuclear and cytoplasmic, in the neurons that were rescued by GFP-bchs-1 (visible in Fig. 7C,G). Interestingly, expression of GFP-bchs-1 in the *bchs* mutant backgrounds resulted in an increase in intensity of Atg5 in vesicles, but a reversal of the Atg5 compartment number accumulation (Fig. 7A,B). GFP-bchs-1 expression caused a strong increase in both Atg8 compartment number and intensity over wild type (Fig. 7E,F). Full multi-cell comparisons of Atg5 and Atg8 antibody stainings of the different genotypes can be seen in montages in Supplementary Figs. 3-6.

**Figure 7.**
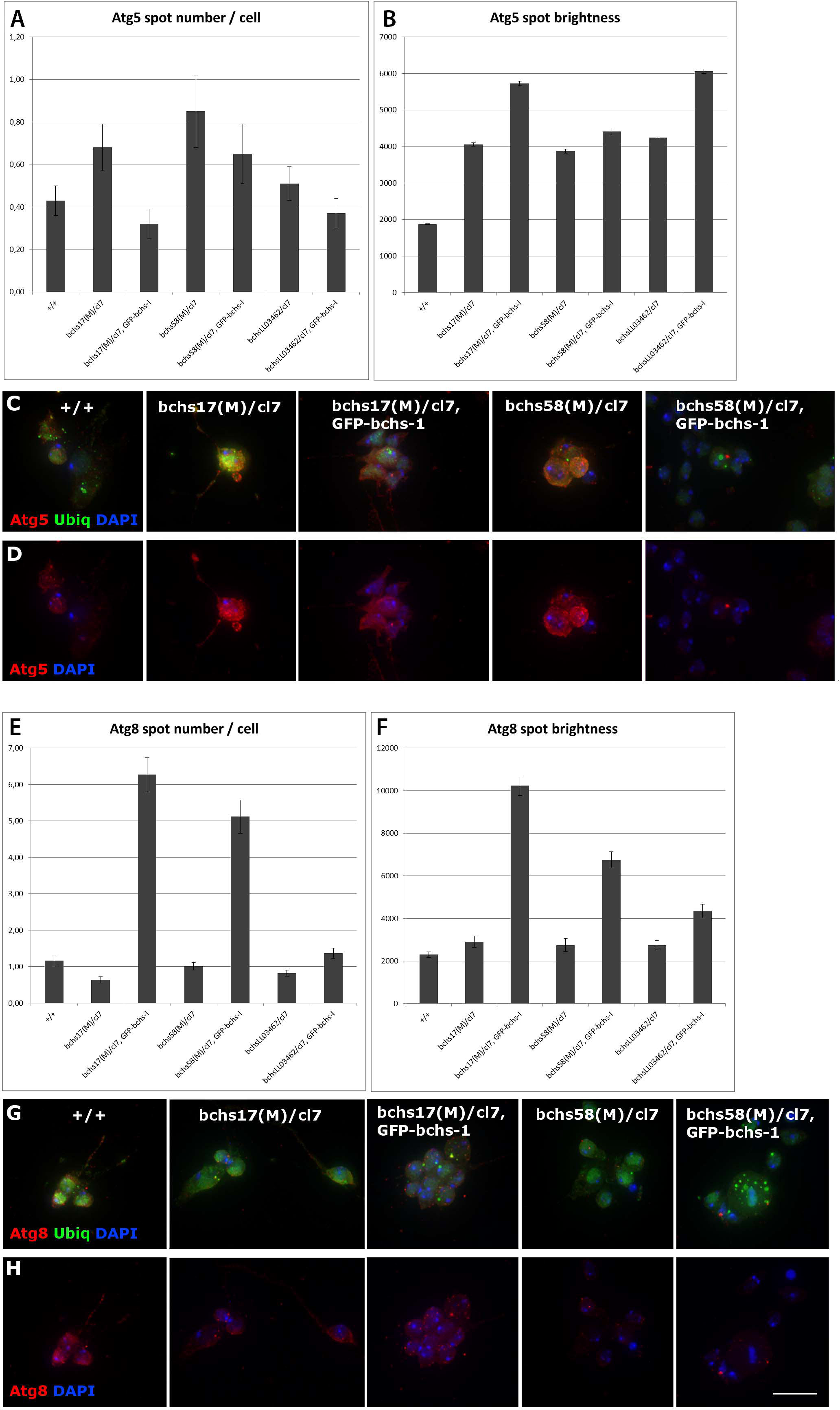
*bchs* mutant primary neurons accumulate immature Atg5-positive autophagic compartments, and expression of the GFP-bchs-1 drives an increase in maturation to Atg8 carrying vesicles. Quantification of Atg5 (red) positive vesicle number **(A)** and brightness **(B**), with examples from individual genotypes **(C, D)**, also labelled with DAPI (blue) to show cell locations and anti-poly-ubiquitin (green). A similar analysis was done for the Atg8 marker on the same genotypes **(E-H)**. Atg5/DAPI and Atg8/DAPI are shown separately to make changes in abundance of the respective marker more visible. Scalebar =10 µm.

We were surprised by the lack of strong changes in Atg8 in mutant neurons, but noted that both Atg5 and Atg8 accumulated detectably in the neurites of *bchs17(M)/cl7* compared to wild-type in neurons that had been allowed to differentiate further by aging for 4 additional days in culture (Fig. 8). This differs from the findings of Filimonenko *et al*^15^ who found that starvation-induced LC3 expression was unaffected in ALFY knockdowns, but this discrepancy may be due to differences in subcellular localization of the accumulations, cell type, or presence vs. absence of autophagy induction. These neurons also showed notably more prominent ubiquitinated aggregates than the younger preparations (Fig. 8, enlarged insets, green). *bchs* neurons with ubiquitin aggregates induced by driving Huntingtin (Htt) Q93 showed no difference from the wild type situation in either Atg5 or Atg8 colocalization with poly-Ubiquitin (Supplementary Fig. 7).

**Figure 8.**
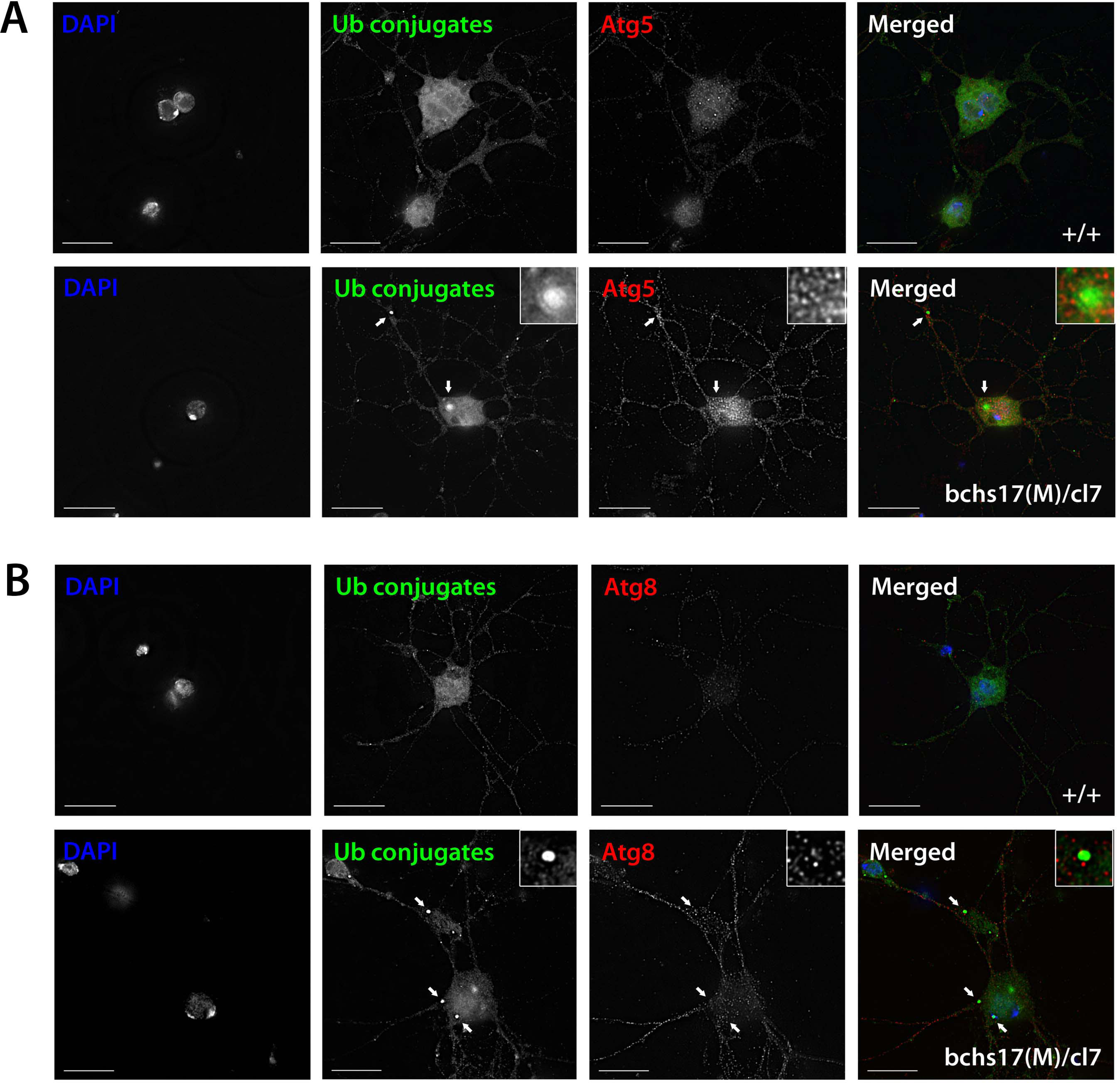
Primary neurons cultured from third instar larval brains of the indicated genotypes, aged 4 days, and labelled with DAPI to show cell location, anti-poly-ubiquitin, **(A)** Atg5 and **(B)** Atg8. The top-right inset is a 5x digital zoom for the indicated region of interest by an arrow showing the deposition of ubiquitinated aggregates. Scalebar =10 µm.

We conclude from the above set of experiments that *bchs* neurons experience an autophagic block leading to a moderate excess of early Atg5 compartments, which can be overcome by expression of the first, full-length Bchs isoform. Indeed, Bchs isoform1 appears to induce super-normal numbers and intensities of Atg8 vesicles (Fig. 7E,F), as well as greater intensities of Atg5. This indicates a likely function of the full-length Bchs protein in maturation of autophagosomal compartments, giving much greater numbers and intensities of mature Atg8-positive compartments when over-expressed. However, aggregates introduced into *bchs*-deficient neurons can still become associated as normal with the autophagic machinery.

### Bchs colocalizes preferentially with Atg5 during selective autophagy and with Atg8 during non-selective autophagy

After autophagy stimulation by nutrient starvation, rapamycin treatment and Htt polyQ93 expression, it was observed generally that the size and number of individual Bchs, Atg5 and Atg8 compartments increased compared to non-treated controls (Figs. 9, 10). However, Bchs was homogeneously distributed throughout the primary neuron and colocalized only ~20% with either marker, RFP-Atg5 or mCherry-Atg8a, under basal and induced autophagy (Figs. 9, 10), which is surprising in light of earlier findings with Alfy^19^. We also note that in cases of overlap between Bchs and the respective markers, the punctae do not completely colocalize with each other (see merged images in figs. 9. 10), suggesting their localization on different compartments or sub-compartments.

**Figure 9.**
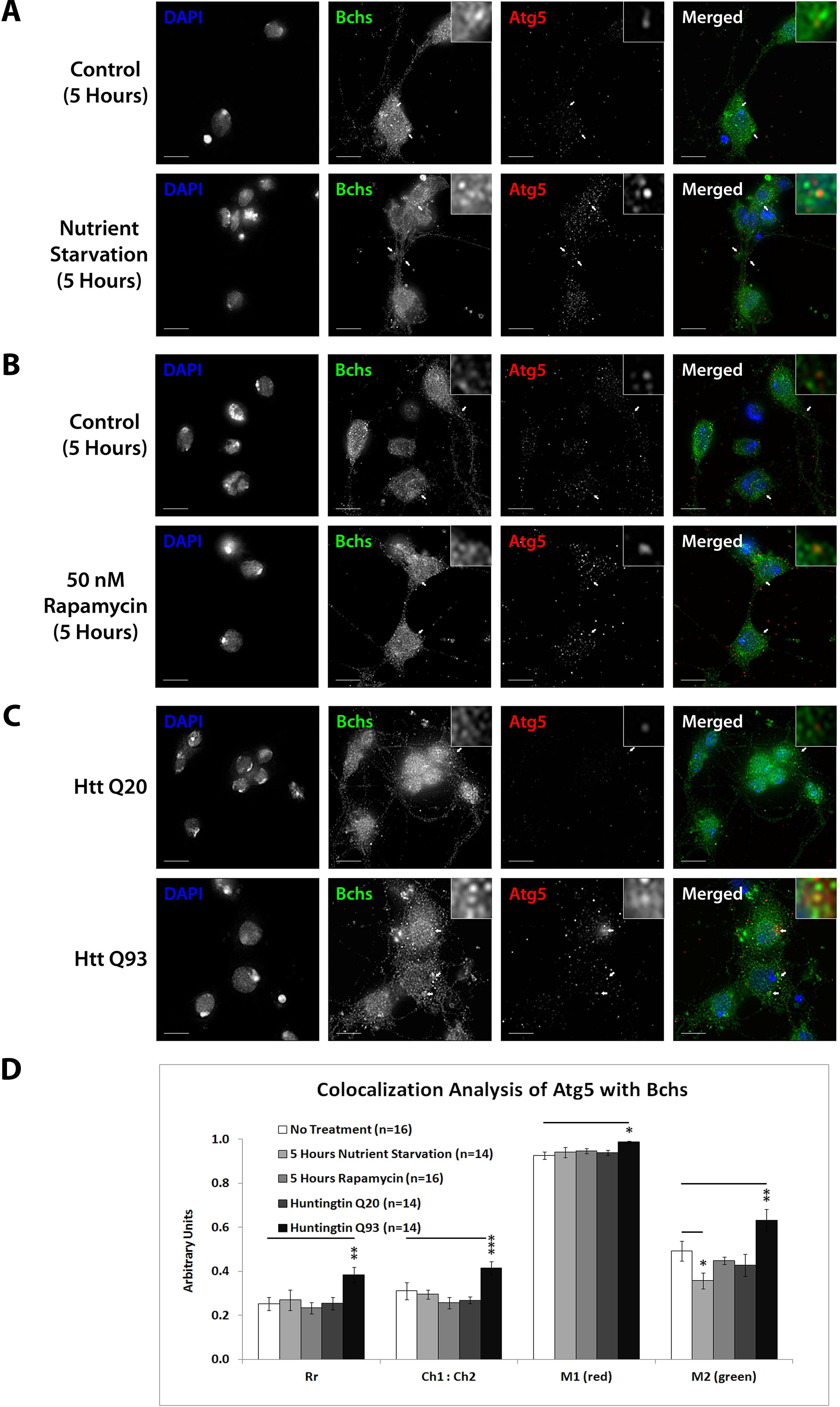
Bchs associates with RFP-Atg5 marked compartments during aggrephagy and dissociates from them during starvation-induced autophagy. Third instar larval primary neurons were immunostained for endogenous Bchs and transgenic RFP-Atg5 using anti-Bchs and anti-DsRed antibodies after different conditions of autophagy induction: **(A)** five hours of nutrient starvation by incubation with serum-free HL-3 buffer, **(B)** five hours of incubation with 50 nM rapamycin in complete medium and **(C)** Htt normal (Q20) and expanded (Q93) polyQ expression. Scalebar = 10 µm. Arrowheads indicate regions of close association between Bchs and RFP-Atg5. The top-right inset shows a 10x magnification of one of the indicated regions. **(D)** Colocalization analysis was performed using ImageJ to obtain Rr (Pearson’s correlation coefficient), Ch1:Ch2 ratio (red:green pixels ratio), M1 (Manders’ colocalization coefficient for Channel 1) and M2 (Manders’ colocalization coefficient for Channel 2). Error bars represent standard error of the mean for n = number of single-slice images. Unpaired Student’s t-test is used for the statistical comparison between treatment and non-treatment groups. * *p* < 0.05, ** *p* < 0.01 and *** *p* < 0.001.

ALFY is localized on nuclear membranes in HeLa cells and only translocates to cytoplasmic structures in the presence of aggregating proteins^15^. In contrast to this, we found Bchs homogeneously distributed throughout primary neurons, as well as in nuclei. Bchs was only observed to be more concentrated on the nuclear membranes occasionally under basal autophagy conditions (Supplementary Fig. 8).

The colocalization of endogenous Bchs with transgenically expressed compartmental markers in response to different stimuli was analyzed using the Intensity Correlation Analysis (ICA) plugin in ImageJ^30,31^. The Manders’ colocalization coefficient for Channel 2 (M2), which expresses the proportion of Bchs-positive (green) pixels overlapping with RFP-Atg5-positive (red) pixels, decreased during nutrient starvation, but not under treatment with rapamycin (Fig. 9A,B,D), another inducer of non-selective autophagy, indicating that the dissociation from Atg5-containing compartments may be a specific response to starvation. In contrast, the expression of Htt Q93 (but not Q20) led to a significant increase in Rr (Pearson’s correlation coefficient), M1 (Manders’ colocalization coefficient for Channel 1) and M2 values with RFP-Atg5 (Fig. 9C,D). The ratio of RFP-Atg5 intensity to endogenous Bchs intensity, expressed by Ch1:Ch2, also increased, although the transgenic RFP-tagged protein is expected to be expressed at constant levels. This may be due to the clearly visible increase in punctate localization of this marker upon autophagic induction, which would be expected to be enhanced by the colocalization algorithm during background subtraction. These results together suggest that Bchs may associate preferentially with early autophagosomal Atg5 compartments during aggrephagy, and at least partially dissociate from them during starvation-induced autophagy.

Nutrient starvation and rapamycin treatment gave a significant increase in the relative quantity of mCherry-Atg8a to Bchs (expressed by the Ch1:Ch2 value), again likely explained by an increase in vesicular localization of the transgenically expressed tagged mCherry-Atg8a protein upon autophagy induction (Fig. 10A,B,D). The amount of Bchs colocalizing with this mCherry-Atg8a (M2) also increased significantly (Fig. 10D). In contrast, Htt Q93 expression did not increase M2 (Fig. 10C,D), as it had with RFP-Atg5 (compare Fig. 9D). These results show that Bchs increases its association with mCherry-Atg8a-tagged compartments after general autophagy induction, but not aggregate induction. A negative control to validate the accuracy of the colocalization algorithm on Bchs and RFP or mCherry immunostaining was performed on *bchs17(M)/cl7* primary neurons not expressing the fluorophore-tagged compartmental markers (Supplementary Fig. 9A).

**Figure 10.**
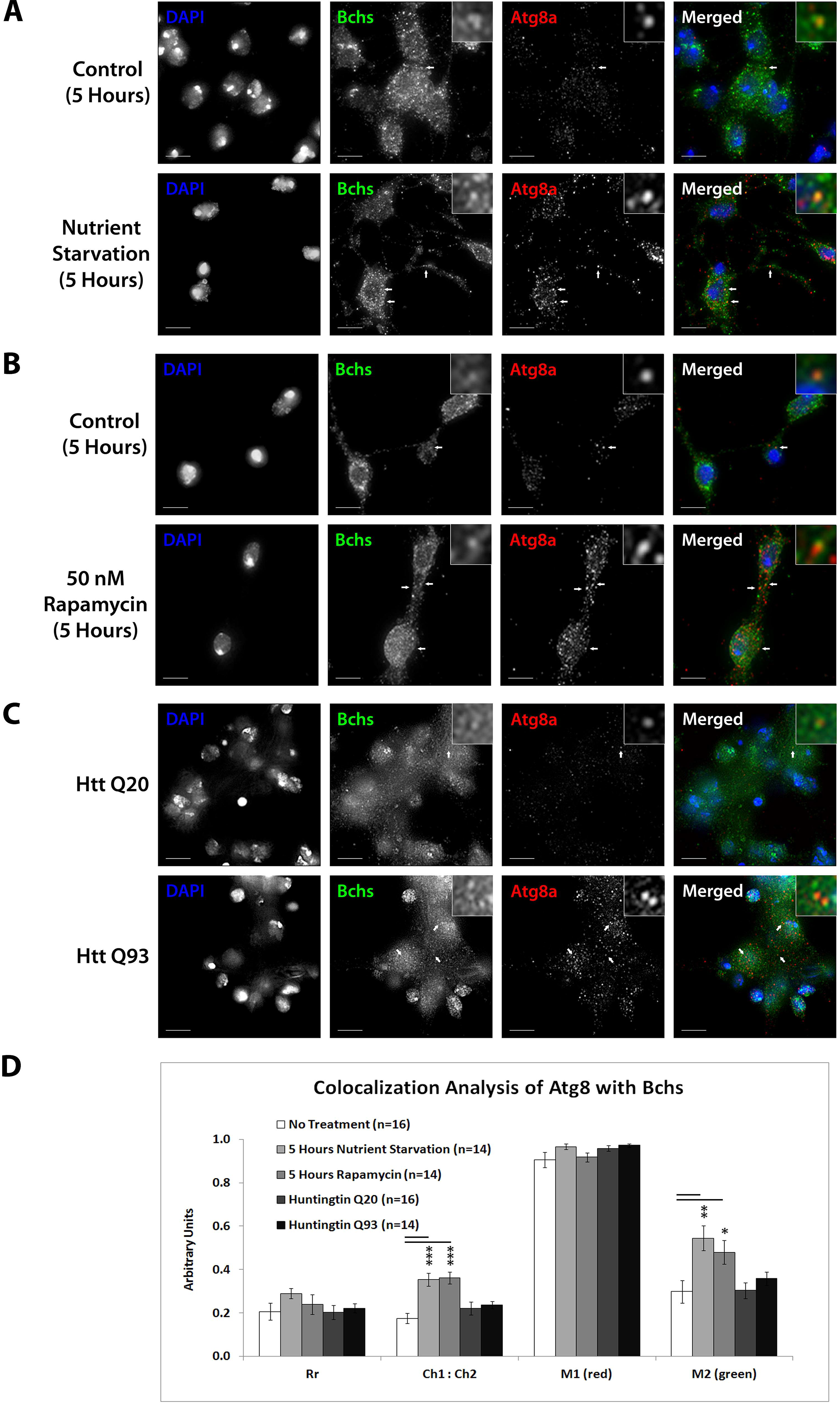
Bchs associates more with mCherry-Atg8a during the induction of non-selective autophagic response. Third instar larval primary neurons were immunostained for endogenous Bchs and transgenic mCherry-Atg8a using anti-Bchs and anti-DsRed antibodies after different conditions of autophagy induction: **(A)** five hours of nutrient starvation by incubation with HL-3 buffer, **(B)** five hours of incubation with 50 nM rapamycin in complete medium and **(C)** Htt normal (Q20) and expanded (Q93) poly-glutamine expression. Scalebar = 10 µm. Arrowheads indicate regions of close association between Bchs and mCherry-Atg8a. The top-right inset shows a 5x magnification of one of the indicated regions. **(D)** Colocalization analysis was performed using ImageJ to obtain Rr (Pearson’s correlation coefficient), Ch1:Ch2 ratio (red:green pixels ratio), M1 (Mander’s colocalization coefficient for Channel 1) and M2 (Mander’s colocalization coefficient for Channel 2). Error bars represent standard error of the mean for n = number of single-slice images. Unpaired Student’s t-test is used for the statistical comparison between treatment and non-treatment groups. * *p* < 0.05, ** *p* < 0.01 and *** *p* < 0.001.

### Selective autophagy results in lowered Bchs association with Rab11-GFP compartments

Rab11 localizes onto recycling endosomes and is required for early endocytic membrane trafficking and recycling^32,33^. Bchs colocalizes partially with Rab11-GFP in *Drosophila* embryonic motorneurons and Rab11 antagonizes Bchs function in synaptogenesis^18,20^. To look at how this interaction is affected by autophagy, the spatial relationship of Bchs with Rab11-GFP was investigated in primary neurons in response to autophagy induction by different conditions. After autophagy induction by all three methods – nutrient starvation, rapamycin treatment or Htt polyQ expression – there was a reduction in colocalization between Bchs and Rab11-GFP as measured by both the Pearson’s correlation coefficient (Rr) and M1 (the proportion of Bchs overlapping with Rab11-GFP), with Htt polyQ giving the strongest reduction (Fig. 11A-D). These data show that Bchs associates less with Rab11 recycling compartments during the induction of selective (aggregate-induced) than under non-selective (starvation or rapamycin-induced) autophagy. However, the number of rab11-carrying compartments appeared not to be affected by autophagy induction. This was tested using a YFP insertion into the endogenous rab11 gene (a kind gift of M. Brankatschk and S. Eaton^34^) in primary neurons treated with rapamycin.

**Figure 11.**
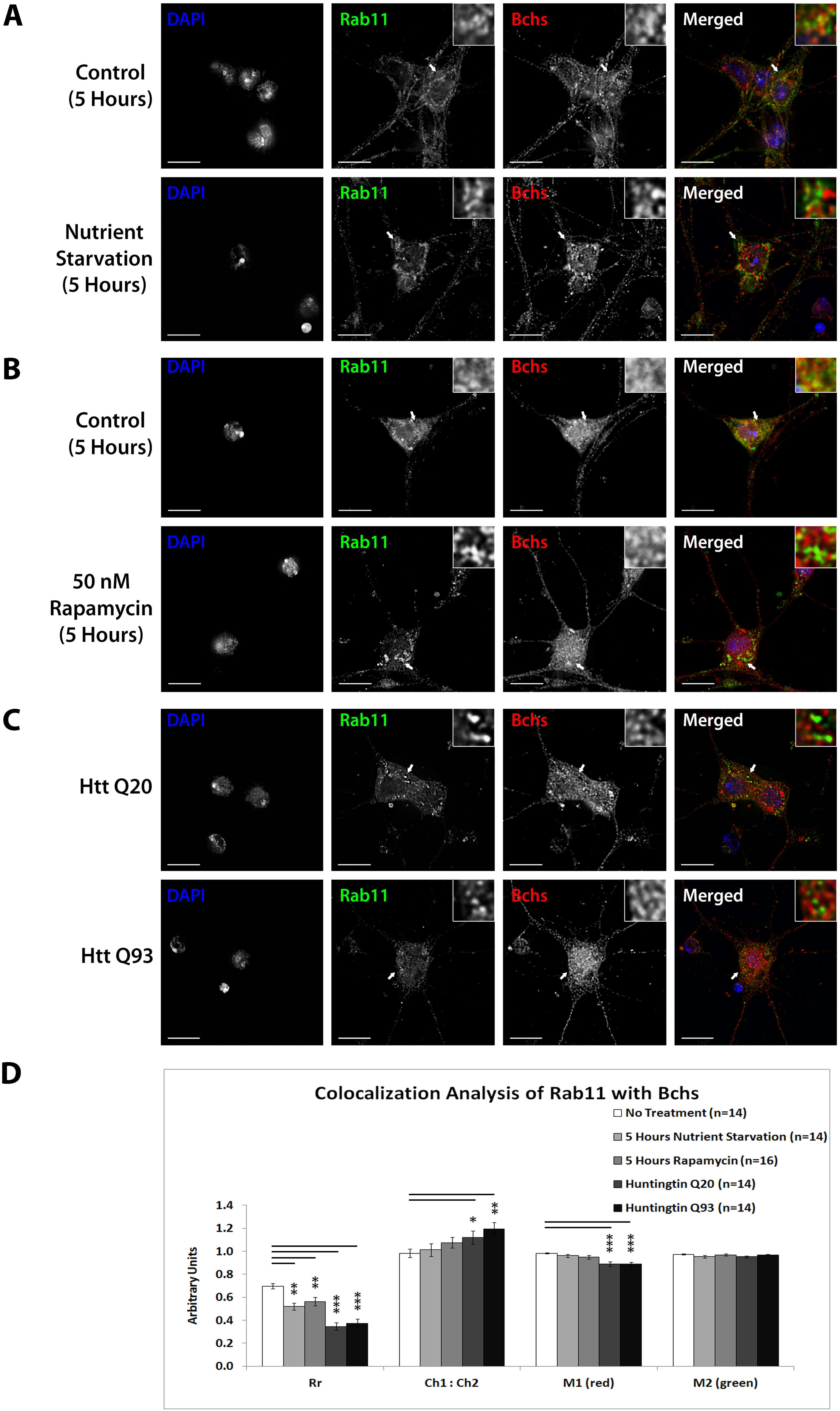
Bchs dissociates more from Rab11-GFP during selective autophagy than non-selective autophagy. Third instar larval primary neurons were immunostained for endogenous Bchs and transgenic Rab11-GFP using anti-Bchs and anti-GFP antibodies after different conditions of autophagy induction: **(A)** five hours of nutrient starvation by incubation with serum-free HL-3 buffer, **(B)** five hours of incubation with 50 nM rapamycin in complete medium and **(C)** Htt normal (Q20) and expanded (Q93) polyQ expression. The top-right inset shows a 5x magnification of one of the indicated regions. Scalebar = 10 µm. **(D)** Colocalization analysis was performed using ImageJ to obtain Rr (Pearson’s correlation coefficient), Ch1:Ch2 ratio (red:green pixels ratio), M1 and M2 (Manders’ colocalization coefficients for channels 1 and 2). Error bars represent standard error of the mean for n = number of single-slice images. Unpaired Student’s t-test is used for the statistical comparison between treatment and non-treatment groups. * *p* < 0.05, ** *p* < 0.01 and *** *p* < 0.001.

For the foregoing colocalization experiments between Bchs and Atg5, Atg8, and Rab11, a negative control to validate the specificity of Bchs and GFP immunostaining was performed on *bchs17(M)/cl7* primary neurons not expressing the fluorophore-tagged compartmental markers (Supplementary Fig. 9B).

### Live imaging of GFP-Bchs with RFP-Atg5 or mCherry-Atg8a during autophagy induction confirms relocalization of Bchs dependent on stimulus

In order to study the dynamics of Bchs interaction with autophagic vesicles, primary neurons cultured from third instar larval brains expressing GFP-bchs-1 and RFP-Atg5 or mCherry-Atg8a were imaged live. GFP-bchs-1 punctae are localized adjacent to RFP-Atg5 punctae occasionally under basal autophagy (Supplementary Fig. 10A; Supplemental movie S1). After nutrient starvation, GFP-Bchs and RFP-Atg5 appeared more distinct (Supplementary Fig. 10B; Supplemental movie S2), in agreement with the reduction in colocalization seen in fixed preparations (Fig. 9D). RFP-Atg5 also re-locates nearer to the nuclear membrane after 40 minutes of starvation (Supplementary Fig. 10B). These structures may correspond to the phagophore assembly sites (PAS), which are thought to form from omegasomes on the ER^35,36^.

After transfection of Htt Q15, occasional GFP-Bchs punctae adjacent to RFP-Atg5 punctae were observed (Supplementary Fig. 10C; Supplemental movie S3), similar to basal conditions. Htt Q128 transfection resulted in observable association of GFP-bchs-1 and RFP-Atg5 in live neurons (Supplementary Fig. 10D; Supplemental movie S4). GFP-bchs-1 in these experiments was distributed mainly homogeneously throughout the cytosol, similarly to endogenous Bchs.

Under basal autophagy conditions, some mCherry-Atg8a punctae were adjacent to GFP-Bchs punctae (Supplementary Fig. 10E, Supplemental movie S5). After two hours of nutrient starvation, however, there was a distinct enlargement of mCherry-Atg8a autophagosomes (Supplementary Fig. 10F; Supplemental movie S6). These “doughnut-like” punctae progressed to a “bean-like” shape resembling the cross-section of vesicular compartments (visible in Supplementary Fig. 10F).

When Htt Q15 or Q128 were expressed, there was little observable coincidence between mCherry-Atg8a and GFP-Bchs in live cells (Supplementary Fig. 10G,H; Supplemental movies S7, S8), similar to basal conditions and corroborating the colocalization analysis (Fig. 10D). mCherry-Atg8a accumulated in a prominent focal swelling along the axon (left arrow in Supplementary Fig. 10H), and an mCherry-Atg8a streak can be seen moving retrograde towards the focal swelling, but does not exit from it towards the cell soma, seeming to indicate a blockage of autophagosomal transport.

## Discussion

In this study, we demonstrate for the first time that neurodegeneration associated with mutations in the proposed aggrephagy adaptor Blue cheese responds positively to autophagy induction, and that newly discovered BEACH-containing isoforms of the protein are necessary for rescue of degeneration by genetic and pharmacological manipulations. This suggests that the BEACH domain, which is known to bind to p62^14,37^, is essential for its role as an autophagy adaptor. Further, we place *bchs* function in an epistatic hierarchy with respect to other components of the autophagy pathway, and show that an intact BEACH isoform is required in order for increased autophagy (e.g. by rapamycin) to be able to rescue neuronal atrophy. This was surprising, since Filimonenko et al found that the WD40/FYVE region alone was responsible for binding Atg5 and could by itself mediate aggregate clearance^15^.

Since motorneuron death caused by strong *bchs* loss of function is not significantly rescued by rapamycin, we propose that Bchs activity functions in autophagy downstream of Atg1^38^. Moreover, our results suggest that neuronal death may not be entirely aggregate-dependent, since the severest motorneuron death is observed in the strong 58(O) allele, but this allele does not show the highest frequency of large terminal aggregates in motorneurons. In fact, the change in localization of Bchs that we observe with different vesicle populations, dependent on the type of autophagy being induced, is consistent with multiple *bchs* functions being involved in neuronal death, in addition to a strict role in aggrephagy^18^.

If autophagy failure is responsible for *bchs* neurodegeneration, then a genetic interaction should be seen between autophagy-modulating conditions and the *bchs* loss of function phenotype, and this was shown for the first time here. Autophagy inhibiting drugs not only exacerbated neuronal death in *bchs* hypomorphs but, importantly, also phenocopied *bchs*, showing that autophagy inhibition alone induces a similar extent of motorneuron death as is seen in *bchs* larvae. However, complete loss of function of *bchs* apparently impairs autophagy to such an extent that it is refractory to further exacerbation by these drugs. We also show that epistatic relationships with Atg7 are strikingly similar to those with autophagy inhibitors and suggest strongly that Bchs acts upstream of Atg7 in the same pathway.

The Atg5/Atg8 studies show an apparent stalling and accumulation of Atg5 compartments in *bchs* mutants, but a greatly increased number and intensity of Atg8 carrying vesicles after overexpression of the long Bchs isoform, suggesting that Bchs may be involved in the progression from earlier to later steps of autophagy, wherein it associates with Atg5-positive phagophores during the induction of aggrephagy (Fig. 12, left). Such a role for Bchs is also consistent with the greatly increased quantity of p62/Ref(2)p marker we documented in an earlier publication^39^. Our epistasis studies additionally show that even in a strong *bchs* loss of function background, an excess of Atg7 can rescue degeneration (see Fig. 4). This suggests that the block in maturation can be overcome by Atg7, although Atg7’s function in conjugation of the Atg12-Atg5-Atg16 complex is nominally upstream of the step toward mature lipidated Atg8-carrying vesicles (Fig. 12, right).

**Figure 12.**
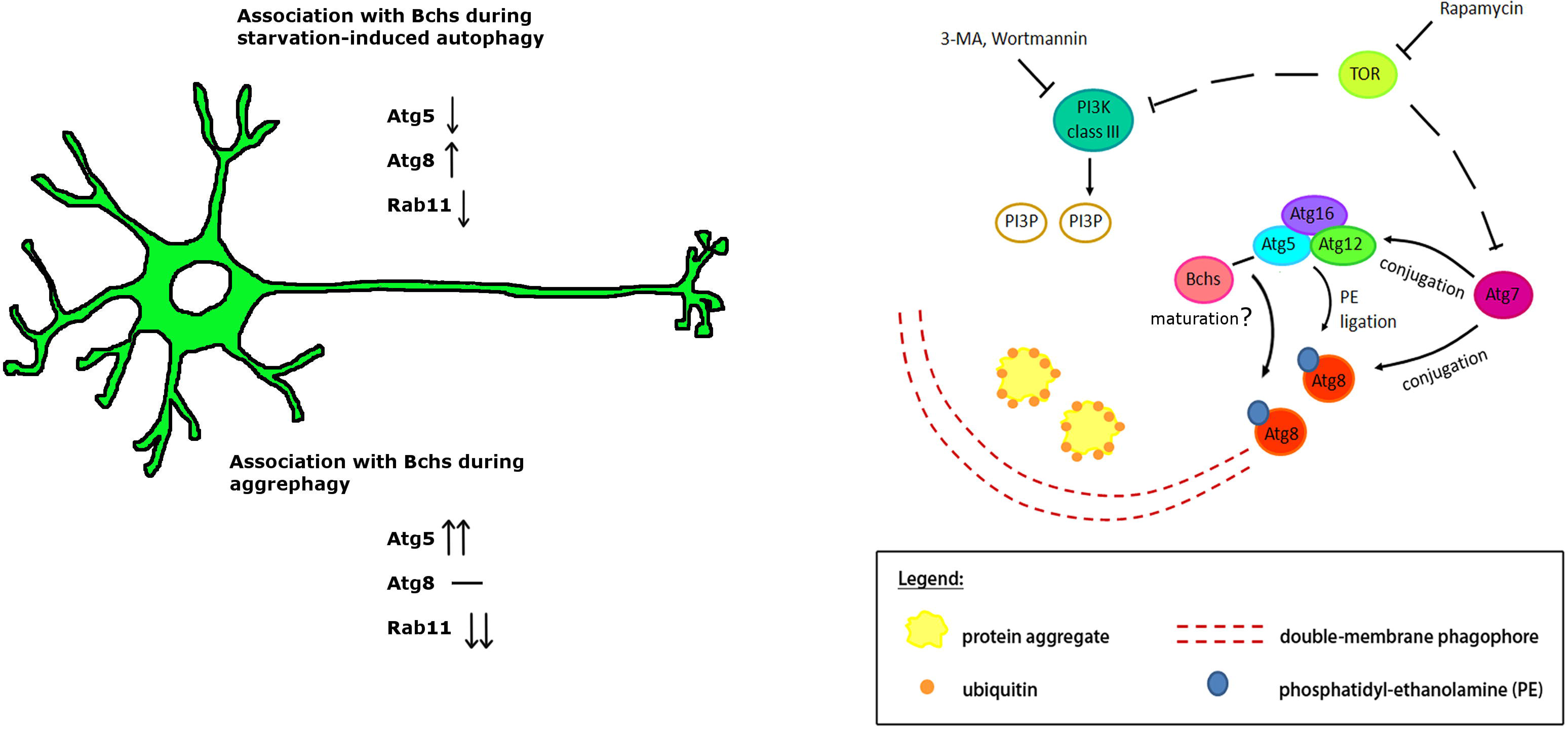
Schematic showing the relationship of Bchs and the autophagy pathway in the clearance of aggregated proteins. Bchs associates preferentially with Atg5 during the induction of aggrephagy and plays a role in maturation of autophagic vesicles to the Atg8-marked stage. Based on epistatic relationships, Bchs acts downstream of the TOR complex and upstream of Atg7 by “priming” target substrates for the autophagy machinery.

In parallel to its increased association with Atg5 during aggrephagy, Bchs dissociates partially from Rab11-positive recycling endosomes after all types of autophagy induction (Fig. 11). Bchs and Rab11 are postulated to perform antagonistic roles^18,20^, with Bchs promoting synaptogenesis while Rab11 inhibits it. Another study reported that autophagy is required for synaptic development^20,40^, an example being Highwire (Hiw) protein, an E3 ubiquitin ligase which suppresses synaptic growth and is degraded by autophagy in synaptic termini. Rab11 may antagonize Bchs through competitive routing of autophagosomal membrane sources under different autophagy stimuli, for example under fed vs. starvation conditions, although a change in rab11 vesicle number, size, or intensity was not apparent after autophagy induction (data not shown)^41,42^.

Taken together, our data suggest that the frequency of association of Bchs with different vesicles depends on whether the autophagy response is selective (aggregate-induced) or non-selective (starvation-induced). We propose that a shift occurs upon the induction of aggrephagy, from membrane recycling to elongation during autophagosome biogenesis, whereupon Bchs may leave recycling endosomes and recruit the Atg12-Atg5-Atg16 E3-like ligase complex for the initiation of autophagosome biogenesis in neuronal termini.

In a previous study, Bchs was reported to be enriched in vesicular structures at synaptic boutons of larval axon terminals^20^. Notably, the initiation of autophagosome biogenesis occurs distally and constitutively in the neurite tips of mouse primary neurons and these then mature as they are transported retrograde in a dynein-dependent manner towards the cell soma^43^. Since aggregates appear prominently in the termini of *bchs* motorneurons, Bchs may be involved in this early step of the autophagy process, which is important in synaptic termini^44^. Interestingly, a recent report showed that fusion between autophagosomes and late endosomes at neuronal termini is required for recruitment of the necessary motor proteins for retrograde transport of the fused compartment, the amphisome^45^. Given the role of BEACH domain proteins in vesicle fusion events, and Bchs’ effects on transport of endo/lysosomes^18^, this seems a likely scenario for a possible function of Bchs.

A number of exocytic SNARE proteins are required at the step of autophagosome biogenesis to promote Atg9 recycling through tubulo-vesicular clustering and membrane fusion at PAS^46^. In addition, the Q-SNARE Syntaxin 17 (Stx17) localizes onto the outer membrane of completed autophagosomes and mediates the fusion with both late endosomes and lysosomes in mammalian cells and *Drosophila*.^47,48^ Intriguingly, the BEACH protein LYST was reported to bind to SNARE proteins^49^, raising the possibility of Bchs interacting with SNAREs to promote vesicle fusion. On the other hand, our finding of increased numbers of Atg8 vesicles after overexpression of the large BEACH isoform is reminiscent of the increased numbers of lysosomes that were seen after LYST overexpression^50^. This is consistent with a similar role in vesicle fission, as LYST and related proteins are thought to perform^51^, whereby BEACH proteins are implicated as accessories in both fission and fusion events of different vesicle populations^52^. Bchs’ possible role in vesicle trafficking and membrane dynamics, in the context of its recently reported genetic interactions with sphingolipid lipases^39^ will be an interesting avenue to investigate in the future.

## Materials and Methods

### *Drosophila* stocks and maintenance

Flies were raised on standard yeast/cornmeal agar food at 25°C. The fly lines used were *bchs58(O), bchs58(M), bchs17(M), Df(2L)clot7, yw; EPgy2 Atg7[EY10058]* (Bloomington Drosophila Stock Center); *w; Atg7[d77]/CyO-GFP*, and *yw; UAS-mCherry-Atg8a* and UAS-Atg1(6A) were kindly provided by Thomas Neufeld. *yw; +; UAS-RFP-Atg5, w; +; w; UAS-Rab11-GFP*, and Htt-polyQ expressing lines *w; UAS-Htt exon1-Q20* and *-Q93* were generous gifts of Katja Köhler, Sean Sweeney, Henry Chang, and Larry Marsh, respectively. The *bchsLL03462* and UAS-Atg1^GS10797^ stocks were obtained from Drosophila Genetic Resource Center in Kyoto Institute of Technology. The eve-Gal4>UAS-CD8-GFP stock (an *even-skipped* driver combined with a membrane-bound GFP marker) was a kind gift of Miki Fujioka.

### Reverse-transcription PCR of *bchs* and *atg7* transcripts and generation of GFP-Bchs

Larval or adult brains were homogenized in 400 µL of TRIzol^®^ reagent (Life Technologies, 15596-026), and total RNA was extracted with chloroform:TRIzol^®^ 1:5. RT PCR was performed using Promega (A5000) M-MLV reverse transcriptase as per manufacturer’s instructions, and PCR was carried out on each cDNA sample with GoTaq^®^ polymerase (Promega, M3005). Gene specific primers used to amplify *bchs* splice isoform 2 were forward = GCAAACAGTTCAGACAATATAC and reverse = AAGATCCTTTATCAGCTGCTTGGC, and splice isoform 3 were forward = GATGGACAGAAAACGATGCTACC and reverse = TTCGCAGGATGAATTTCTCGTG. *atg7* specific primers were: forward = TCCGCAGACGGATTGATCTC and reverse = TGAACATCGATGACAGCCTTG; PCR primers for *rp49* were: forward = AGTCGGATCGATATGCTAAG and reverse = AGTAAACGCGGTTCTGCATG. Full length *bchs* isoform 1 was cloned into pUAST vector containing a GFP tag at the 5’ end. Transgenic animals were generated by Best Gene (Chino Hills, CA). Further details of the cloning can be provided on request.

### Motorneuron viability assay

Third instar larvae were dissected, fixed 30 min in 4% paraformaldehyde (Sigma Aldrich, P6148), blocked 1 h in 5% bovine serum albumin (BSA) (PAA Laboratories, K41-001) in 0.1% Triton X-100 PBS (PBT), and incubated overnight at 4°C with primary antibodies: 1:500 anti-GFP (Clontech Laboratories, 632377), 1:10 1D4 anti-fasciclin II and 1:20 22C10 anti-futsch (Developmental Studies Hybridoma Bank), and washed in PBT. Secondary antibodies were 1:800 Cy2-conjugated goat anti-rabbit and 1:800 Cy3-conjugated goat anti-mouse (Jackson ImmunoResearch Laboratories, 111-225-144 and 115-165-146) in PBT. Motorneuron viability was scored by inspection. RP2 survival was scored as a percentage over total hemisegments and repeated in triplicate. Chi-square statistical test was performed.

### Drug feeding

Embryos were collected every 3 hours on apple juice agar plates, and incubated at 25°C for 24 h; hatching 1^st^ instar larvae were transferred into 2.0 mL Eppendorf tubes with food containing 0.05% (v/v) ethanol (vehicle control), 1 µM rapamycin (AG Scientific), 0.2 or 2 µM Wortmannin (AG Scientific; R-1018) or 5 mM 3-methyladenine (Calbiochem^®^, Merck Millipore, 189490), and dissected as third instar larvae.

### Quantitative size analysis of ubiquitinated aggregates in larval NMJs

Aggregates were detected with 1:1000 mouse anti-poly-ubiquitin (clone FK2, Enzo Life Sciences, BML-PW8810) and 1:800 Cy5-goat anti-mouse and 1:500 Cy2-goat anti-horse radish peroxidase (Jackson ImmunoResearch Laboratories, 115-175-146 and 123-095-021). Image stacks were acquired under 40x objective lens with a DeltaVision OMX^®^ microscope (Applied Precision) and deconvolved with softWoRx^®^ 5.0. To measure aggregate area, ImageJ particle analysis was used.^32^ The measurement scale (in microns) was defined on the image projection, adjusted to a segmentation threshold of 70, and converted to a binary image. A region of interest (ROI) was marked and ‘Analyze Particles’ used to measure the area of particles within the ROI. Particle size was categorized into 0 - 1 µm^2^, 1.1 - 10 µm^2^ and 10.1 - 50 µm^2^. Percentages of NMJs with aggregates in each of the three groups were calculated for each experimental set, which was repeated in triplicate, and a Chi-square statistical test was performed.

### Primary neuron culture from third instar larval brains, and imaging of GFP-bchs-1

The protocol was modified from Kraft and co-workers.^53^ Larvae were washed with 90% ethanol three times and sterile water twice to remove debris and contaminants. Four third instar larval brains were extracted in Shields and Sang, bacto-peptone and yeast extract (Sigma Aldrich, S8398) medium containing penicillin-streptomycin and antibiotics-antimycotics (PAA Laboratories, P11-002), washed with sterile hemolymph-like 3 (HL3) saline (70 mM NaCl, 115 mM sucrose, 5 mM trehalose, 5 mM KCl, 20 mM MgCl_2_, 1 mM CaCl_2_, 10 mM NaHCO_3_, 5 mM HEPES, pH 7.4) three times, and incubated with 0.5 mg/mL collagenase type 1 (Sigma Aldrich, C1639) in HL3 1 h at room temperature. Brain tissue was then washed three times with complete medium with 10% heat-inactivated fetal bovine serum (HyClone, Thermo Fisher Scientific, SH30070.03), 20 µg/mL of bovine pancreas insulin (Sigma Aldrich, I1882), penicillin-streptomycin and antibiotics-antimycotics. Brains were dissociated into a final volume of 150 µL as a single cell suspension by trituration and then aliquoted onto 22 mm^2^ acid-washed cover-slips coated with 30 µg/mL of mouse laminin (BD Biosciences, 354232) and 167 µg/mL of concanavalin A (Sigma Aldrich, C0412) in 35 mm culture dishes and allowed to attach ~12 h. 900 µL of complete medium was added to the culture dish and incubated between 1 - 4 d.

For the colocalization in Fig. 2, primary neurons from larval offspring of elav-Gal4>mCherry-Atg8a or elav-Gal4>UAS-RFP-Atg5 crossed with UAS-GFP-bchs-1, 2, or 3 were live-imaged at 60x magnification on the DeltaVision microscope as above, using Z-stack acquisition of 200 nm optical slices with interleaving green and red channels at 150 msec exposures. 100 image stacks for each colocalization were examined at each Z-level, and where vesicular localization of the Bchs isoform was detectable, the individual vesicle was then visually inspected in red/green image pairs for complete, non-, or partial overlap of Bchs with mCherry-Atg8a or RFP-Atg5. The colocalization algorithm used below was not applied here because of a high cytoplasmic expression level of the GFP-Bchs isoforms. Each vesicle expressing Bchs was only counted once. Standard error was calculated by averaging the percentages in each of the three categories (“yes”, “no”, or “partial”) over the data separated into three equal batches of images.

### Immunocytochemistry and image analysis of endogenous Atg5 or Atg8 compartments in *bchs* primary neurons

*bchs* and wild-type primary neurons were fixed with 4% (w/v) paraformaldehyde (Sigma Aldrich, P6148) in 100 mM PBS for 15 minutes at room temperature and blocked with 5% (v/v) normal goat serum (Life Technologies, 50-062Z) in 0.05% (v/v) Triton X-100 PBS for 1 h. Antibody staining was performed as above with 1:1000 mouse anti-poly-ubiquitin (clone FK2, Enzo Life Sciences, BML-PW8810) and 1:500 rabbit anti-drAtg5 (Novus Biologicals, NB110-74818) or 1:500 rabbit anti-drAtg8a (a kind gift of Katja Köhler). Samples were incubated with 1 µg/mL of 4’,6-diamidino-2-phenylindole (DAPI), dihydrochloride (Life Technologies, D1306) in PBS for 3 min, washed briefly with PBS, then water, and imaged at 100x magnification for aged primary neurons, and 60x for 1-day old primary neurons.

For the image analysis in Fig. 7, images of primary neuron cultures were acquired as Z-stacks, with the step of 200nm in between slices, using the same exposure, light intensity and filter settings for every condition. Collected stacks were then deconvolved as above, and converted into 16-bit TIFF stacks in FIJI (http://fiji.sc) using LOCI BioFormats plug-in^54^. Each stack was then Z-projected using the maximum intensity method. Images were inspected under DAPI channel and only single cells, or small aggregates (<5 cells) with clearly separated cell nuclei were used for further spot counting. Because of varying spot intensity relative to cell body, and variable background signal, spots were counted manually, after setting the same display range of values. The cell number N used for calculations varied between conditions (N_average_ ~100), but was not less than N_min_=50. The average spot number per cell and the standard error of the mean (SEM) were calculated. To measure spot characteristics, projected images were masked using the Maximum Entropy threshold method^55^. Brightness and size distribution and SEM of selected spots were measured using the Analyze Particles FIJI command.

### Quantitative colocalization analysis of Bchs with different compartmental markers

Autophagy was induced in primary neurons by five hours of nutrient starvation with HL3, five hours of 50 nM rapamycin in complete medium or Huntingtin Q93 expression. Controls (basal autophagy) were incubated in complete medium. Antibody staining was performed as above. For mCherry-Atg8a or RFP-Atg5, primary antibodies were 1:500 rabbit anti-Bchs and 1:100 mouse anti-DsRed (BD Pharmingen™, 551814). For Rab11-GFP, primary antibodies were rabbit anti-Bchs and 1:1000 mouse anti-GFP (Clontech Laboratories, 632375). Colocalization analysis was done with the ImageJ plugin ‘Intensity Correlation Analysis’ after channel splitting and background subtraction^30^. Rr (Pearson’s correlation coefficient), Ch1:Ch2 ratios, M1 and M2 (Manders’ colocalization coefficient for channel 1 and 2) were tabulated for each image. Unpaired Student’s t-test was used to calculate *p* values in between treated and control groups.

### Time-lapse imaging of GFP-Bchs with RFP-Atg5 or mCherry-Atg8a in primary neurons

Primary neurons expressing *GFP-Bchs* and *RFP-Atg5* or *mCherry-Atg8a* transgenes via elav-Gal4 were cultured on 35 mm glass bottom dishes (World Precision Instruments, FD35-100) coated as described above. For basal autophagy, cells were incubated in complete culture medium at room temperature and single focal-plane images were acquired for GFP-Bchs: green channel = 50% transmission, 800 ms; RFP-Atg5: red channel = 50% transmission, 800 ms; and mCherry-Atg8a: red channel = 32% transmission, 600 ms, every 10 min over 4 h, and deconvolved. For nutrient starvation, neurons were incubated in HL3 saline at room temperature and immediately imaged as described. For transfection of Huntingtin 15Q or 128Q, 3 µL of FuGENE^®^ HD transfection reagent (Promega, E2311) was used to couple 1 µg of pCINeoHtt1955.15Q.wt or pCINeoHtt1955.128Q.wt plasmid DNA (kind gift of Anat Yanai and Mahmoud Pouladi) respectively in a 3:1 ratio in a final volume of 50 µL complete culture medium without penicillin/streptomycin and antibiotics/antimycotics for 10 min at room temperature, and added to the cells with 1 mL of complete culture medium without penicillin/streptomycin and antibiotics/antimycotics for 48 hours (without medium change).

## Acknowledgements

The authors sincerely thank final year project students in the School of Biological Sciences, Liyanah Mohd Zaffre, Hong Hwee Lim, Bruno Yip, and Weixin Xu, for data contributing to the live imaging analysis with GFP-bchs, the Wortmannin and rapamycin studies. Funding was provided by the Singapore Biomedical Research Council grant #09/1/22/19/610 and the Ministry of Education of Singapore grant #MOE2013-T1-002-229, a Bundesministerium für Bildung und Forschung (BMBF) Singapore-Germany cooperation grant, and the Eleonore Trefftz Programme for visiting Women Professors at TU Dresden.

